# Processing of the Same Narrative Stimuli Elicits Common Functional Connectivity Dynamics Between Individuals

**DOI:** 10.1101/2022.11.25.517964

**Authors:** Başak Türker, Laouen Belloli, Adrian M. Owen, Lorina Naci, Jacobo D. Sitt

**Affiliations:** Sorbonne Université, Institut du Cerveau - Paris Brain Institute - ICM, Inserm, CNRS, Paris 75013, France; Instituto de Ciencias de la Computacion, CONICET-UBA, Buenos Aires, Argentina; The Western Institute for Neuroscience, Western Interdisciplinary Research Building, University of Western Ontario, London, ON N6A 5B7, Canada; Trinity College Institute of Neuroscience, School of Psychology, Trinity College Dublin, Lloyd Building, Dublin, Ireland

## Abstract

It has been suggested that the richness of conscious experience can be directly linked to the richness of brain state repertories. Brain states change depending on our environment and activities we engage in by taking both external and internally derived information into account. It has been shown that high-level sensory stimulation changes local brain activity and induces neural synchrony across participants. However, the dynamic interplay of cognitive processes that underlie moment-to-moment information processing remains poorly understood. Using naturalistic movies as an ecological laboratory model of the real world, here we assess how the processing of complex naturalistic stimuli alters the dynamics of brain networks’ interactions, and how these in turn support information processing. Participants underwent fMRI recordings during movie watching, scrambled movie watching, and rest. Measuring phase-synchrony between different brain networks, we computed whole-brain connectivity patterns. We showed that specific connectivity patterns were associated with each experimental condition. We found a higher synchronization of brain patterns across participants during movie watching compared to resting state and scrambled movie conditions. Moreover, synchronization increased during the most engaging parts of the movie. The synchronization dynamics across participants were associated with suspense; more suspenseful scenes induced higher synchronization. These results suggest that processing of the same high-level information elicits common neural dynamics among individuals and that whole-brain functional connectivity tracks variations in the processed information and the subjective experience.

## INTRODUCTION

The content of our conscious experience changes depending on the environment and ongoing task. Both external and internal information are processed and integrated to give rise to our conscious experiences. The dynamic interplay of cognitive processes that underlie our moment-to-moment experience of the world remains poorly understood. Using naturalistic movies as an ecological laboratory model of the real world, previous studies have shown that audio-visual clips influence brain activity in a similar manner across individuals, linked to their common conscious experiences and wholistic understanding. Movie watching can synchronize brain activity in the cortex^1–6^, elicit time-resolved correlations between pairs of regions^7^, and give rise to consistent whole-brain activations across participants^8^. Moreover, studies have shown that the quality of encoding of the movie’s content is correlated with inter-subject synchronization during movie watching^9,10^. However, the descriptions of local activations offer only a limited summary of the dynamic processes that give rise to coherent understanding over time.

In recent years, dynamic descriptions of brain activity have gained prominence as they might better account for the participant’s mental state. Focusing on how different brain networks interact over time, rather than the classic description of local activity, could provide a better understanding of conscious processing. The interaction between brain regions has been widely investigated using static functional connectivity^11–14^ computed over the entire fMRI scan (for a comprehensive review see^15^). More recently, dynamic functional connectivity measures came into use, revealing transient brain states that vary in time^16–20^, reflecting cognitive processes at any given moment^21,22^. It has been suggested that the richness of conscious experience can be directly linked to the richness of brain state repertories. Indeed, individuals who lack consciousness present brain states that are less diverse, with fewer long-range interactions and no anticorrelation between brain areas^23–25^. Furthermore, active interventions (i.e. deep brain stimulation) that aim to induce recovery from a state of unconsciousness also increase brain state repertory richness^26^.

A recent study representing brain network interactions in a latent space found increased overall similarity between participants who actively tried to understand a scrambled movie^27^. However, the movie features that drive inter-subject synchronization of brains states are poorly understood. Given the evolving nature of movie plots, high-level cognitive processing of the movie varies significantly over time. Therefore, unravelling the relationship between similar brain state dynamics across participants and the feature dynamics of a particular narrative is necessary for understanding how brain processes give rise to movie comprehension over time.

To address this gap, we examine how the processing of plot-driven naturalistic movies dynamically shapes brain state repertoire, as measured via dynamic brain networks’ interactions. We characterize whole-brain connectivity patterns that emerge during resting state, movie watching, and scrambled movie watching in healthy participants. We show that certain brain patterns are more frequent during movie-watching whereas some others are more frequent in non-movie conditions. Moreover, by assessing temporal mean and dynamic (time-resolved) synchronization of the connectivity patterns between participants, we show that narrative stimuli induce higher inter-subject synchronization, especially during suspenseful scenes. Altogether, these results suggest that processing of the same narrative stimuli elicits common functional connectivity configurations between individuals, and the dynamics of these brain states track variations in the high-level properties of the processed information.

## RESULTS

We investigated how high-level sensory information processing influences ongoing brain activity emerging from the coordination of different brain regions. 15 participants underwent fMRI recordings during movie watching and rest. A second group of 12 participants watched the same movie but scrambled to prevent them from understanding the plot while still viewing every scene. Using Hilbert transform and k-means clustering, we computed whole-brain connectivity patterns for each fMRI volume in each condition (Figure 1). The clustering procedure has resulted in four distinct connectivity patterns. While Pattern 1, 2 and 3 showed both positive and negative correlations between different brain networks, Pattern 4 lacked long-range connectivity (Figure 2A).

**Figure 1.**
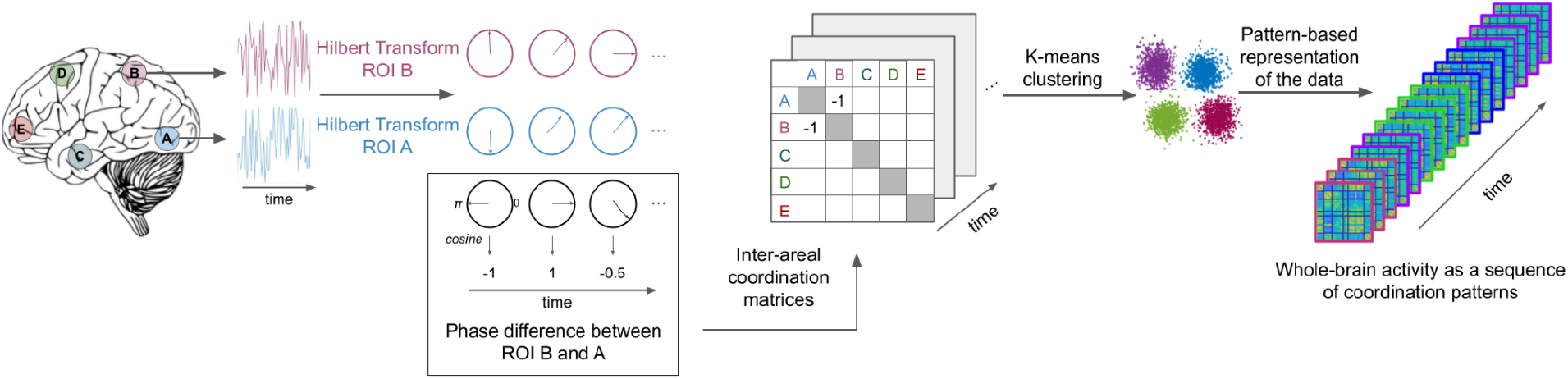
Summary of the method for inter-areal connectivity pattern computation. Following the preprocessing of the fMRI data, BOLD signal time-series were extracted from 42 regions of interest (ROIs) belonging to 6 different networks (visual, auditory, saliency, default mode, fronto-parietal, and motor). The Hilbert transform was applied to all time-series in order to extract the instantaneous phase at each fMRI volume (pink and blue circles in the figure). Phase-differences were computed between each ROI pair (in this example between ROI A and B) at each time point (black circles in the figure). Using cosine similarity, we ranged the phase synchronies between −1 and 1; −1 indicating a complete phase opposition and 1 indicating a complete phase coherence between the two ROIs. Then, phase synchrony values were used to create a 42 by 42 inter-areal coordination matrix for each time-point (fMRI volume). Using k-means clustering, we classified the coordination matrices into 4 *prototypical patterns*. Finally, the fMRI data of each participant was expressed as a sequence of these 4 patterns.

**Figure 2.**
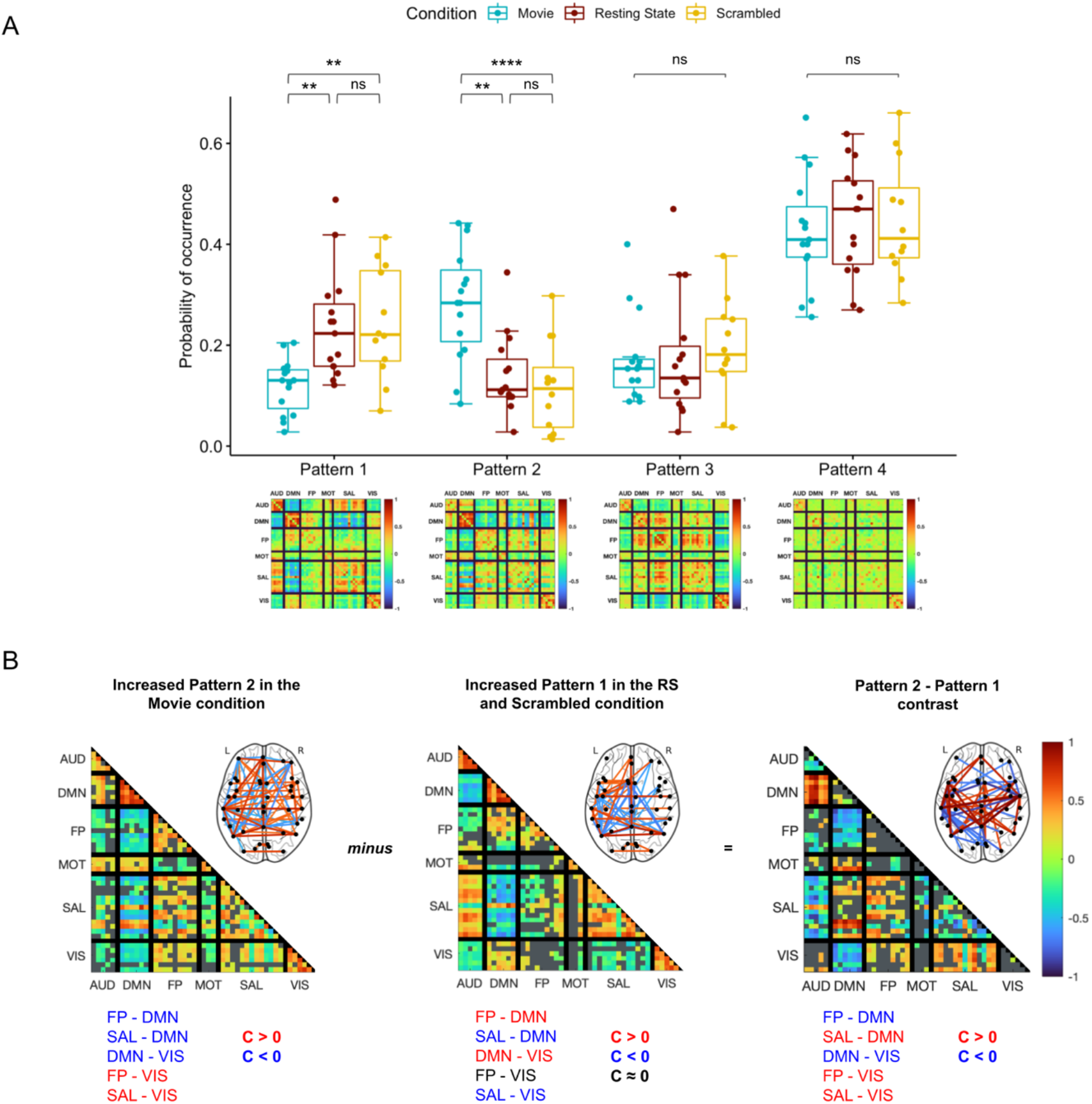
Condition specific variation of the whole-brain connectivity patterns. (A) Occurrence probability of the 4 patterns during movie watching (blue), rest (brown) and scrambled movie watching (yellow). The occurrence probability of Patterns 3 and 4 did not differ in the three conditions. However, Patterns 1 and 2 showed condition-specific modulation: while Pattern 2 showed increased occurrence during movie watching, Pattern 1 was more frequent in scrambled movie watching and resting-state conditions. Stars indicate statistical significance after correction for multiple comparisons using the Bonferroni procedure. (B) Description of the differences between Pattern 2 (more frequent in movie condition) and Pattern 1 (more frequent in non-movie conditions). Blue shades indicate negative coherence (C<0)) whereas red shades signify positive coherence (C>0) after correction for multiple comparisons using the Benjamini–Hochberg procedure (Sign test; *p* < 0.001). Gray color marks coherence that is not significantly different than zero. Note that brain templates show top 5% coherence for visualisation purposes. Pattern 2 (left), which was more frequent in the movie condition, showed negative phase coherence between the default-ode network (DMN) and fronto-parietal (FP), saliency (SAL) and visual (VIS) networks while the visual network had positive coherence with fronto-parietal (FP) and saliency (SAL) networks. Pattern 1 (center), which had more occurrence in non-movie conditions, was characterized by positive coherence between default-mode network (DMN) and fronto-parietal (FP) and visual (VIS) networks and negative coherence between saliency network (SAL) and visual (VIS) and default-mode networks. The contrast between patterns (Pattern 2 - Pattern 1; right) revealed more negative coherence between default-mode network (DMN) and fronto-parietal (FP) and visual (VIS) networks in Pattern 2 (increased with movie) than Pattern 1 (increased with non-movie conditions). Moreover, saliency (SAL) and visual (VIS) networks showed more positive coherence in Pattern 2 compared to Pattern 1. Gray color indicates p values greater than 0.001 after FDR correction in a mixed linear model.

First, we hypothesized that some whole-brain connectivity patterns would be associated with a particular experimental condition and would change expression frequency accordingly. A linear model revealed a Pattern*Condition interaction (*F(6)* = 7.17, *p* < 0.0001). Pattern 2 showed increased occurrence during movie watching (median: 28.4%) compared to resting state (median: 11.2%, *t(156)* = 3.94, *p* = 0.0004, after Bonferroni correction), and scrambled movie watching (median: 11.4%, *t(156)* = 4.77, *p* < 0.0001). By contrast, Pattern 1 was less frequent during movie watching (median: 13.0%) compared to resting state (median: 22.3%, *t(156)* = −3.48, *p* = 0.0019) and scrambled movie (22.1%, *t(156)* = −3.38, *p* = 0.0027) (Figure 2A). We didn’t find any difference between the resting state and scrambled movie watching for Patterns 1 (*t(156)* = −0.099, *p* = 1.00) and Pattern 2 (*t(156)* = 1.05, *p* = 0.88). Patterns 3 and 4 had similar occurrence probabilities in all conditions.

Given the movie-specific modulation of Patterns 1 and 2, we investigated how they differed in their inter-ROI coherence profiles. Pattern 2 -more frequent during movie watching– showed positive coherence between the saliency network (SAL), fronto-parietal (FP) and visual (VIS) networks, and negative coherence between the default-mode network (DMN) and fronto-parietal (FP), saliency (SAL) and visual (VIS) networks (Figure 2B, left panel). On the other hand, Pattern 1 -more frequent in the non-movie conditions-exhibited negative coherence between SAL - DMN and SAL - VIS, and positive coherence between DMN - FP and DMN - VIS (Figure 2B, middle panel). The contrast between Pattern 2 and Pattern 1 (Figure 2B, right panel) revealed that Pattern 2 contained more negative coherence between DMN - FP and DMN - VIS, and more positive coherence between FP - VIS and SAL - VIS compared to Pattern 1.

Next, we hypothesized that functional connectivity dynamics would be similar across participants during movie watching when the same narrative drives brain activity. We computed an inter-subject similarity index (ISI) that allows us to assess the inter-subject co-occurrence of the patterns in the three experimental conditions. Since we were describing the brain activity with only four patterns, co-occurrence of patterns could arise by chance, even though participants were under different conditions and their brain activity was completely independent. The ISI indicates how much more co-occurrence participants have compared to chance-level co-occurrence. As expected, we found ISI values around zero during rest (Wilcoxon signed rank test after Bonferroni correction: *V* = 35, *p* = 0.51) and scrambled movie watching (*V* = 64, *p* = 0.16), indicating that the co-occurrence of patterns during those conditions was at chance level. We observed a significant increase in the ISI during movie watching (median ISI = 0.05, S.E. = 0.005, *V* = 120, *p* = 0.0002)) compared to rest (median SI = 0, S.E. = 0.003, *t(39)* = 9.56, *p* < 0.0001, after Bonferroni correction) and scrambled movie watching (median SI = 0.01, S.E. = 0.004, *t(39)* = 6.74, *p* < 0.0001), indicating higher co-occurrence when participants watched the movie.

Moreover, we predicted that the co-occurrence of the patterns would be more important during the most engaging parts of the movie, during which the movie plot captures the attention. We took advantage of the suspenseful nature of the movie and utilized a pre-existing dataset where a third group of 15 participants had rated the suspense of the scenes on an 8-item scale outside of the scanner^1^. We noted higher suspense values in the second half of the movie compared to the first half (mean: 1.3 vs 0.9, Wilcoxon rank-sum rest: *z* = 2.43, *p* = 0.01). As predicted, ISI values were higher only in the second half of the movie (*F*(2) = 44.34, *p* < 0.0001) and did not differ between conditions during the first half of the recordings (*F*(2) = 1.73, *p* = 0.2) (Figure S1). The same was true for the patterns’ occurrence probabilities in the three conditions, which only differed in the second half of the recordings (Figure S2). For a finer assessment of inter-subject brain state synchronization, we used entropy as a measure of instantaneous pattern co-occurrence among participants at each time point. Lower entropy values indicate higher co-occurrence between participants. We observed a negative relationship between the average suspense ratings and the entropy values: scenes with higher suspense were followed by a decrease in entropy, indicating a higher co-occurrence across participants (Figure 3B). The relationship between the suspense and entropy was further confirmed by a significant Spearman’s correlation (*rho* = −0.25, *p* = 0.0002), taking a 6 second fMRI response lag (3 TRs) relative to the time of suspense rating into account (Figure 3C). This relationship was not found in the resting state (*rho* = −0.08, *p* = 0.25) and scrambled movie conditions (*rho* = 0.09, *p* = 0.21) (Figure S3).

**Figure 3.**
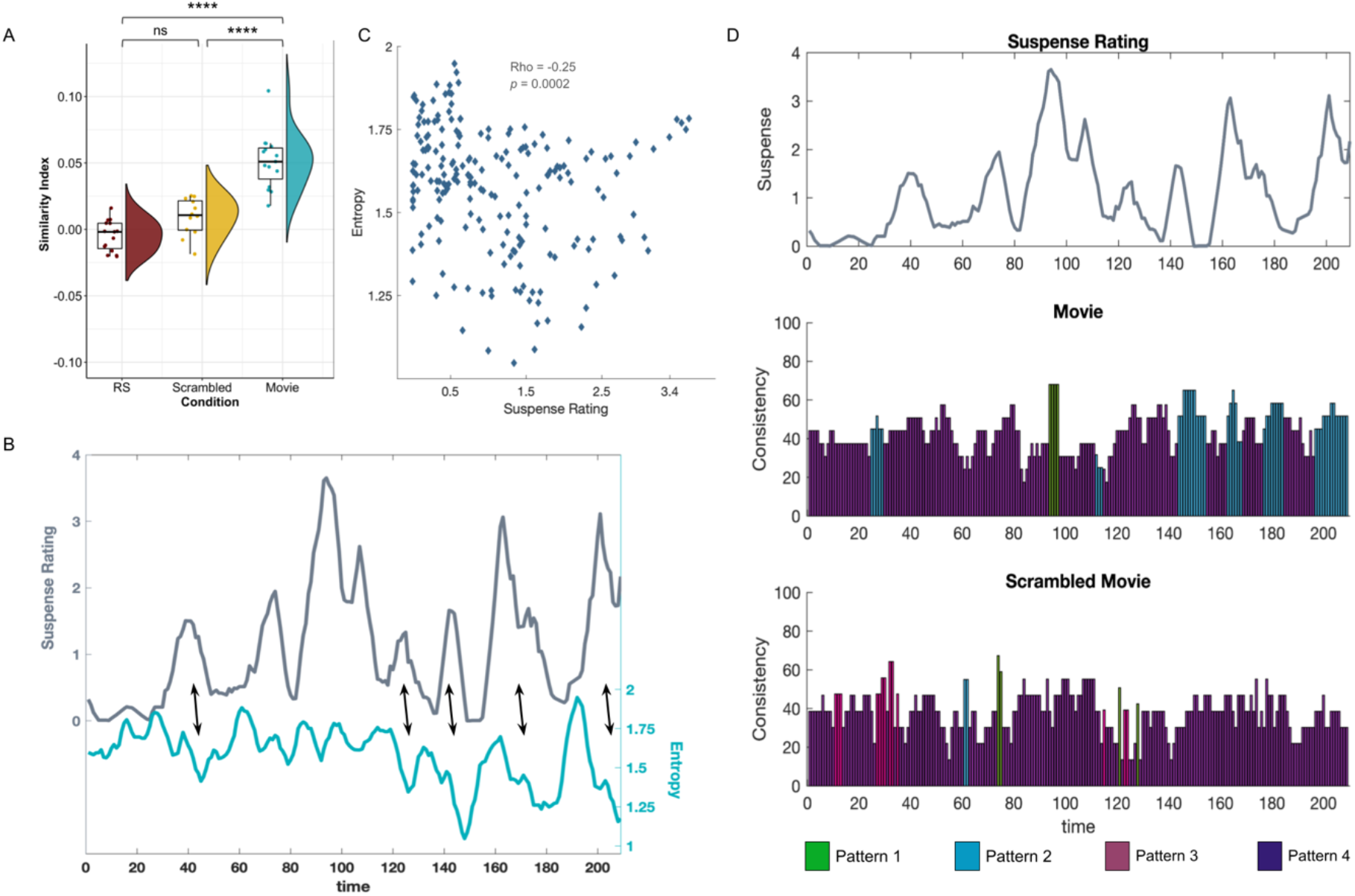
Movie watching induces inter-subject synchronization by increasing the co-occurrence of the movie-related whole-brain connectivity pattern. (A) Average inter-subject similarity index of each participant in resting-state (RS), scrambled movie, and movie watching conditions. Similarity indices were not different than zero in non-movie conditions, indicating a chance-level co-occurrence of patterns among participants throughout the whole duration. On the other hand, participants in the movie condition showed higher similarity index values, indicating increased co-occurence of the patterns compared to chance. Stars denote statistical significance after correction for multiple comparisons using Benjamini–Hochberg procedure. (B) Suspenseful scenes increase pattern co-occurrence during movie watching. The dark gray line shows the variations in the average suspense rating. Entropy (light blue line) of the pattern distribution among participants at each time point is used as an instantaneous co-occurrence measure. Lower entropy values indicate higher co-occurrence. Note the negative relationship between the suspense rating and the entropy values: scenes with higher suspense ratings are followed by a decrease in entropy and thus an increased co-occurence of the patterns among participants. (C) Scatter plot of the average suspense rating and the subsequent entropy values (6 seconds after the suspense rating). We found a significant negative correlation between the two measures (*rho* = −0.25; *p* = 0.002). (D) Instantaneous co-occurrence during suspenseful scenes is associated with the movie-specific pattern. Temporal consistency of the whole-brain connectivity patterns in the movie (middle panel) and scrambled movie (bottom panel) watching condition. Variations in the average suspense rating can be found in the top panel. The dominant pattern at a given time point is indicated by a color code. The y-axis shows the percentage of across-participant consistency. The overall occurrence probability of each pattern in the conditions (Figure 2A) is subtracted from the instantaneous consistency in order to assess consistency exceeding random occurrence. Note the higher consistency values in the movie condition compared to the scrambled movie condition. Pattern 2 (blue) was the dominant pattern during increased suspensefulness, reaching up to 65% excess consistency during movie watching.

Finally, we investigated whether this increased co-occurrence during suspenseful scenes was due to a specific connectivity pattern, or if all patterns contributed similarly to the increase. Following a previously published method^8^, we computed the most frequently expressed pattern at each time window during movie watching and scrambled movie conditions and assessed their consistency across participants (Figure 3D). We subtracted the *time-averaged* occurrence probabilities of each pattern during the movie and scrambled movie watching (Figure 2A) from the consistency level in these conditions to reveal consistency exceeding mean session occurrence. Overall, we found higher consistency values during movie watching compared to scrambled movie watching (mean: 44.1 vs 38.6, Wilcoxon rank-sum test: *z* = 3.65, *p* = 0.0003). Furthermore, Pattern 2, which showed increased occurrence during movie watching, was the dominant pattern during the moments of suspense in the second half of the movie, reaching up to 65% excess consistency.

In sum, our results revealed that (i) dynamic connectivity patterns exhibit condition-specific modulation in their occurrence frequency. Pattern 2, which was characterized by positive coherence between the VIS and FP networks and negative coherence between the DMN and VIS, FP, and SAL networks, shows increased occurrence probability during movie watching compared to non-movie watching conditions; (ii) participants show synchronization in their dynamic functional connectivity profiles during movie watching; (iii) this synchronization is significantly associated with the narrative of the movie, and suspenseful moments are associated with higher synchronization; and (iv) Pattern 2 contributed the most to the synchronization in suspenseful moments. Altogether these results suggest that processing of the same high-level information elicits similar functional connectivity dynamics that reflect and track changes in the properties of the processed information.

## DISCUSSION

In the current study, we investigate how the processing of an engaging audio-visual stimulus alters ongoing brain activity and induces common functional connectivity dynamics across viewers. Until now, most studies investigating inter-subject synchronization during naturalistic stimuli viewing focused on local static activations^1,2,4–6,9,10^. Here we adopt a global and dynamic view of neural activity and show that watching a plot-driven and engaging movie shapes the interaction of brain networks and synchronizes the network dynamics across-individuals. Importantly, we also show that the neural synchronization across participants is not stable but rather fluctuates over time. Indeed, the dynamics of the network interactions vary with the suspense of the scenes, resulting in a higher inter-subject synchronization when participants are immersed in the movie.

In light of these results, we suggest that suspense ratings may be used as a proxy for the allocation of attention. Suspenseful scenes capture attention and therefore will enhance the processing of the scenes, resulting in a higher inter-subject synchronization during these scenes. Compatible with our results, several studies have found that increased suspense in the narrative induces higher activity in the ventral attention network^28,29^. Our results go beyond this finding and show that increased attention in turn, modulates the ongoing processing similarly across individuals.

The two patterns that showed condition-specific modulation present both positive and negative phase coherence between different brain regions that coordinate according to the ongoing task (Figure 2B). For example, positive coherence between visual and FP networks and negative coherence between DMN and FP networks during movie watching suggest that while attention networks coordinate with sensory areas, they decouple from the regions that are implicated in internal thought generation^30,31^. These results are consistent with previous studies reporting a juxtaposition between DM and FP networks during movie watching^32^ and during switches between internal and external awareness^33^, and furthermore show that this decoupling occurs simultaneously with the coupling of sensory and attention networks. Interestingly, studies show that patterns which exhibit both positive and negative coherence between different brain regions are predominant in healthy populations and diminish in unconscious states^23–26^. Our results add to this literature by showing that, in healthy participants, these rich coherence patterns can be divided into similar yet distinct sub-patterns reflecting the cognitive processes implicated in the ongoing task.

One could ask why scrambled movie watching and resting-state induce similar connectivity patterns although the two conditions differ drastically, especially at the sensory level. This might be due to the fact that participants could not follow the plot in the scrambled movie condition and therefore, may instead, have directed their attention to their self-generated thoughts, as they did during resting-state scans^31^. Since the whole-brain patterns capture the global brain state and not region-specific activations, they are likely more sensitive to participants’ ongoing mental activity than detailed sensory-driven processes, and thus, pick up on the similarities between the mental states elicited by these apparently different conditions.

In summary, our results show that movie watching induces synchronization of static and dynamic functional connectivity patterns across participants. This synchronization effect can be used to probe conscious processing in different states such as sleep, anesthesia, or disorders of consciousness (DoC). For example, by investigating the extent to which a DoC patient shows mental processes synchronized with those of healthy controls, we could assess their ability to process external information. Such measures could help the diagnosis procedure, especially for patients exhibiting cognitive-motor dissociation^34^, who are often wrongly diagnosed as being in a vegetative state due to a lack of behavioral responses^35^. These findings have implications for developing tools that probing intact capacities in a range of behaviorally unresponsive populations.

## METHOD

### Participants and Procedure

In this study, we used a previously published dataset^1^ in which 27 participants underwent functional MRI recordings. During the acquisition, 15 participants (18-40 years; 7 males) watched an 8-minute black and white movie clip taken from a TV show entitled “Alfred Hitchcock Presents—Bang! You’re Dead”. The same participants also went through a resting-state scan. A second group of 12 participants (18-30 years; 4 males) watched the same movie but in a scrambled order. In this condition, the movie was cut into 1-second segments and shuffled, allowing participants to see all the scenes without understanding the plot of the movie. To evaluate how suspense varied in the movie, a third group of 15 participants (19–29 years; 5 males) watched the movie clip outside of the scanner and were asked to rate how suspenseful every 2-second segment of the movie was using an 8-point scale. All participants were right-handed native English speakers without any neurological or psychiatric disorders. They signed a consent form prior to the experiment and were remunerated for the participation. This research was approved by the local ethics board of the Western University. Further information on the participants or experimental procedure can be found in Naci et al. (2014)^1^.

### MRI acquisition parameters

MRI data were acquired on a 3T Siemens Tim Trio System. T2*-weighted whole-brain images were recorded during resting state (256 volumes), movie (246 volumes) and scrambled movie (238) watching with a gradient-echo EPI sequence (33 slices, slice thickness: 3 mm, interslice gap of 25%, TR/TE: 2000 ms/30 ms, voxel size: 3 × 3 × 3 mm, flip angle: 75°). A mirror box allowed participants to see the movie that was presented on a projection screen behind the scanner. Noise cancellation headphones (Sensimetrics, S14) were also used for sound delivery. An anatomical volume was also acquired using T1-weighted MPRAGE sequence in the same acquisition sessions (154 slices, matrix size: 240 × 256 × 192, TE: 4.25 ms, voxel size: 1 × 1 × 1 mm, flip angle: 9°).

#### fMRI preprocessing

Raw MRI data were preprocessed and denoised using CONN functional connectivity toolbox^35^ implemented in MATLAB (The MathWorks). The first 5 volumes were discarded to ensure stable magnetization. The preprocessing procedure included realignment, slice-time correction, outlier detection, segmentation, normalization into the MNI152 space (Montreal Neurological Institute), and spatial smoothing using a Gaussian kernel of 6-mm full width at half-maximum. For outlier correction, images with more than 0.3 mm framewise displacement in one of the z, y, z directions, more than 0.02 rad rotational displacement, or global mean intensity exceeding 3 standard deviations were included as nuisance regressors in the generalized linear model (GLM). White matter and cerebrospinal fluid masks were also included as nuisance parameters in the GLMs. Average time-series from 42 regions of interest were extracted after applying a 0.008 to 0.09 Hz band-pass filter to the signal. Regions of interest were defined as 10 mm-diameter spheres around the given MNI coordinates (Table S1).

#### Time-varying functional connectivity patterns

All computations were performed in MATLAB. Following the preprocessing, extracted ROI time-series were represented in the complex space using their analytic representation that composed by the original signal (real part) and the Hilbert transform of the signal (imaginary part). The instantaneous phase is computed as the inverse tangent of the ratio of the imaginary and real components and wrapped into the [−*π*,*π*] interval using the angle function implemented in MATLAB. This allowed us to have a time-series of instantaneous phases for each ROI. Note that to avoid edge artefacts, the first and last 9 time points have been discarded from the time series. Then, phase differences between each ROI pair (861 in total) were calculated at each time-point using cosine similarity. Thus, the whole-brain connectivity configuration at each time point could be represented as an observation in a 861-dimensional space. We concatenated data from all sessions (42 in total: 15 resting states, 15 movies and 12 scrambled movies) and applied k-means clustering (k = 3, 4, 5, 6, and 7) with 1000 repetitions using *Manhattan distance* in order to uncover ‘prototypical’ connectivity configurations that were recurrent in all experimental conditions. The silhouette method revealed that 4 clusters provided the best classification. Connectivity configuration at each time point was then labeled with one of the 4 cluster centroids to which the configuration belonged. Finally, participants’ brain activity during the scans was expressed as a sequence of the 4 centroids (42 by 42 phase coherence matrix).

#### Inter-subject similarity index

We computed an Inter-subject Similarity Index (ISI) in order to assess the resemblance between the participants’ pattern sequences using the following formula

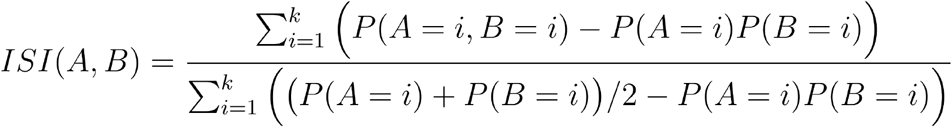

where P(A = *i*) and P(B = *i*) are the occurrence probabilities of Pattern *i (*among total number of k patterns, in this study k = 4) in participants A and B respectively. P(A = *i*, B = *i*) corresponds to the joint probability of exhibiting Pattern *i* by participants A and B at a given time-point. This index measures the co-occurrence of brain patterns between two participants (A and B) while correcting for the overall occurrence probability of each connectivity pattern. Positive values of ISI indicate higher co-occurrence of patterns compared to chance (SI = 0). The synchronization level of each participant is defined as their average SI with all the other participants of their group.

#### Suspense rating

15 participants watched the movie outside of the scanner. The movie was cut into 2 seconds segments. After viewing each segment, participants rated the suspense level of the segment on an 8-point scale ranging from not suspenseful at all to maximal suspense. A moving average of 7 ratings (14 seconds) was computed in order to bring out the changes in the movie plot and remove the rapid variations within the scenes. Average suspense ratings can be found in Figure 3B.

#### Instantaneous co-occurrence

We used entropy as a measure of instantaneous pattern co-occurrence between participants. Entropy H(X) of the connectivity patterns at a given time point is computed with:

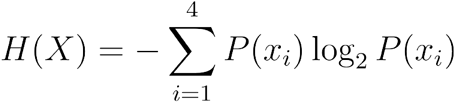

Where P(*x_i_*) is the occurrence probability of pattern *i* among the participants at a given moment. Since we have four distinct patterns, the maximum entropy is equal to 2, indicating a uniform distribution of the pattern probabilities (¼ of the participants exhibited each pattern). Conversely, lower entropy values indicate that majority of the participants had the same pattern at the given time point. This co-occurrence measure is not pattern specific: increased co-occurrence of any patterns would decrease the entropy values. In order to assess the relationship between the suspense variations and the instantaneous co-occurrence among participants, we took the moving average of 7 entropy values as with the suspense ratings.

#### Temporal pattern consistency

Following a previously published method^8^, we assessed which patterns contributed the most to the inter-subject co-occurrence of the patterns. This method consists in counting the number of participants having a given pattern at least once in a time window and in determining the pattern that manifested the most. Here we used a sliding window of 7 consecutive time points to construct the time windows similarly to all our other analyses. The dominant patterns during movie watching and scrambled movie watching can be found in Figure 3D in the center and bottom panels respectively. The percentage of consistency indicates the percentage of participants having the dominant pattern at a time point. In order to exclude consistency that might appear by chance given the natural occurrence probabilities of the patterns, we subtracted the occurrence percentage of a pattern during the movie and scrambled movie-watching scans from the percentage of consistency of this pattern at a given time-point in these conditions.

#### Statistical analyses

Occurrence probabilities of the patterns and changes in the similarity index across conditions are assessed with linear models using lme4^37^, car^38^, and emmans^39^ packages in R^40^. Wilcoxon rank-sum tests on the suspense ratings and the temporal consistency values, as well as the Spearman’s correlation between the suspense values and the entropy were performed on MATLAB (The MathWorks). All tests are corrected for multiple comparisons using the Bonferroni procedure except for the sign-tests on Pattern coherence levels which were corrected using the Benjamini–Hochberg procedure.

## ACKNOWLEDGEMENTS

This study was funded by the European Union (ERA PerMed JTC2019 “PerBrain”, grant to JDS) and the Canada Excellence Research Chairs (CERC) program (Grant No. 215063 to AMO). BT received a PhD grant from French Ministry of Higher Education and Ecole Normale Supérieur. AMO is a Fellow of the CIFAR Brain, Mind, and Consciousness program. Authors would like to thank Garance Merholz for her feedback on the manuscript and Esteban Munoz Musat for helpful discussions.

---

**Figure S1.**
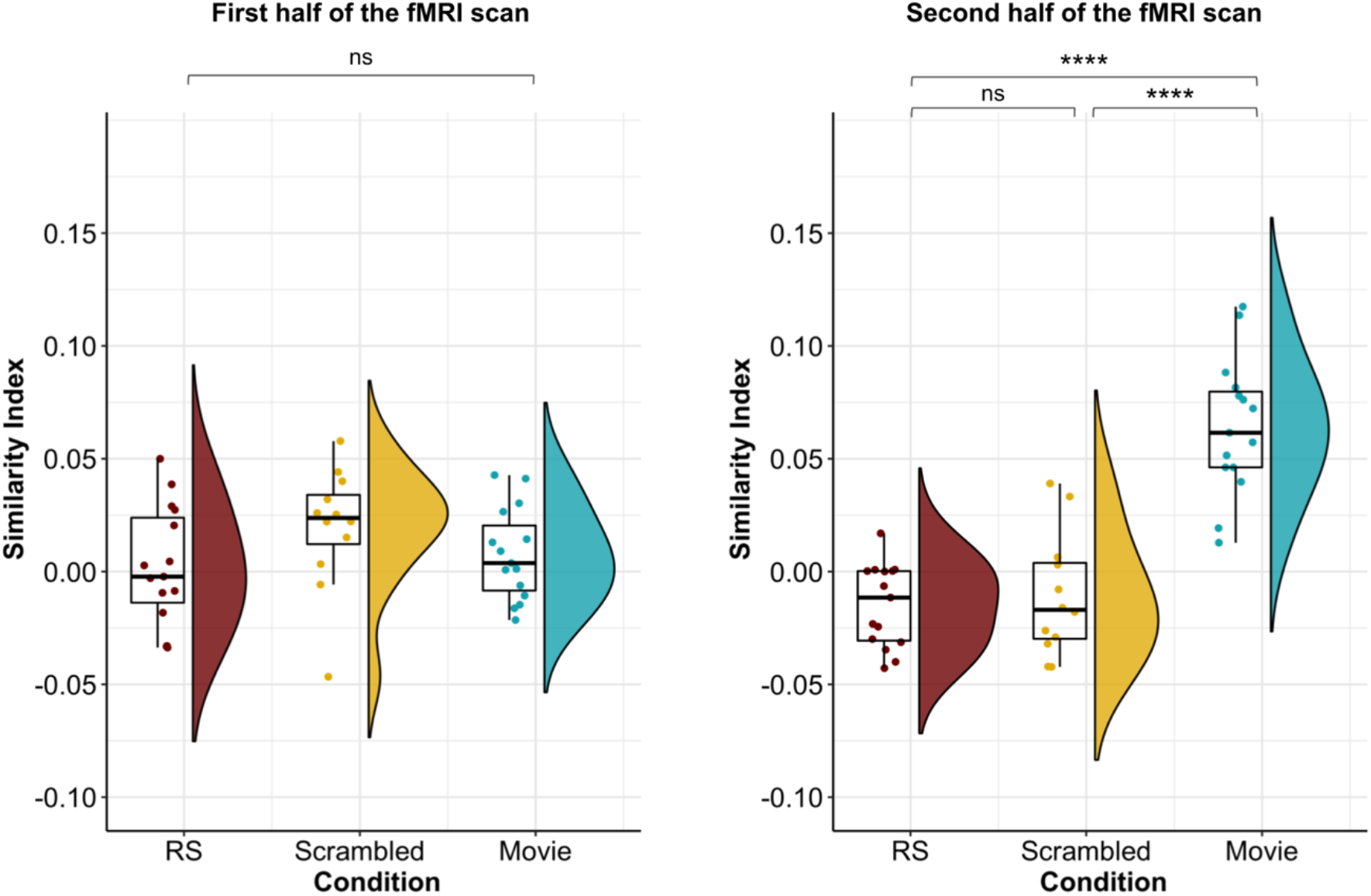
Increased inter-subject similarity index only during the second half of the movie. Average similarity index (ISI) of each participant computed over the first (left) and second (right) half of the fMRI scan during resting-state, scrambled movie watching, and movie watching conditions. While SI did not differ between condition during the first half of the fMRI scan (*F*(2) = 1.73, *p* = 0.19), SI during movie watching was significantly higher compared to resting-state (*t*(39) = 8.49, *p* < .0001) and scrambled movie (*t*(39) = 7.6, *p* < .0001) conditions. SI during resting-state and scrambled movie conditions were not statistically different (*t*(39) = 0.41, *p* = 1). Each dot represents a participant.

**Figure S2.**
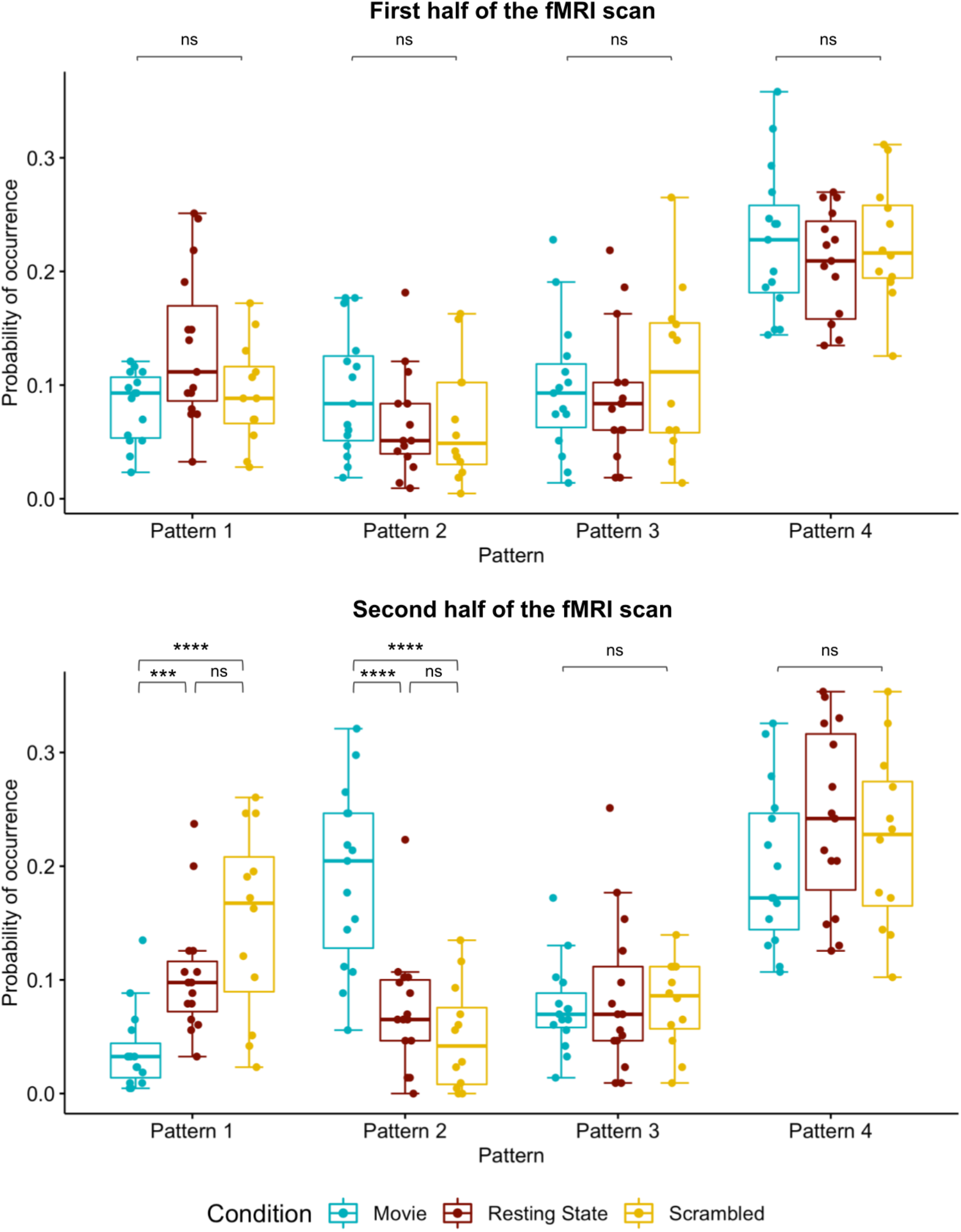
The occurrence probability of the patterns differs between conditions only during the second half of the fMRI scan. The occurrence probability of the patterns during the first (left) and the second (right) half of the fMRI recording in movie watching, resting-state and scrambled movie watching conditions. While no significant interactions were found between the condition and the pattern probabilities during the first half of the scan (*F*(6) = 1.81, *p* = 0.1), a significant interaction was observed in the second half (*F*(2) = 13.00, *p* < .0001). Pattern 2 showed increased probability during movie watching compared to resting-state (*t*(156) = 5.09, *p* < .0001) and scrambled movie watching (*t*(156) = 6.43, *p* < .0001) and Pattern 1 was more frequent during scrambled movie watching (*t*(156) = 5.20, *p* < .0001) and resting-state (*t*(156) = 3.82, *p* = .0006) compared to movie watching. Patterns 3 and 4 did not differ between conditions.

**Figure S3.**
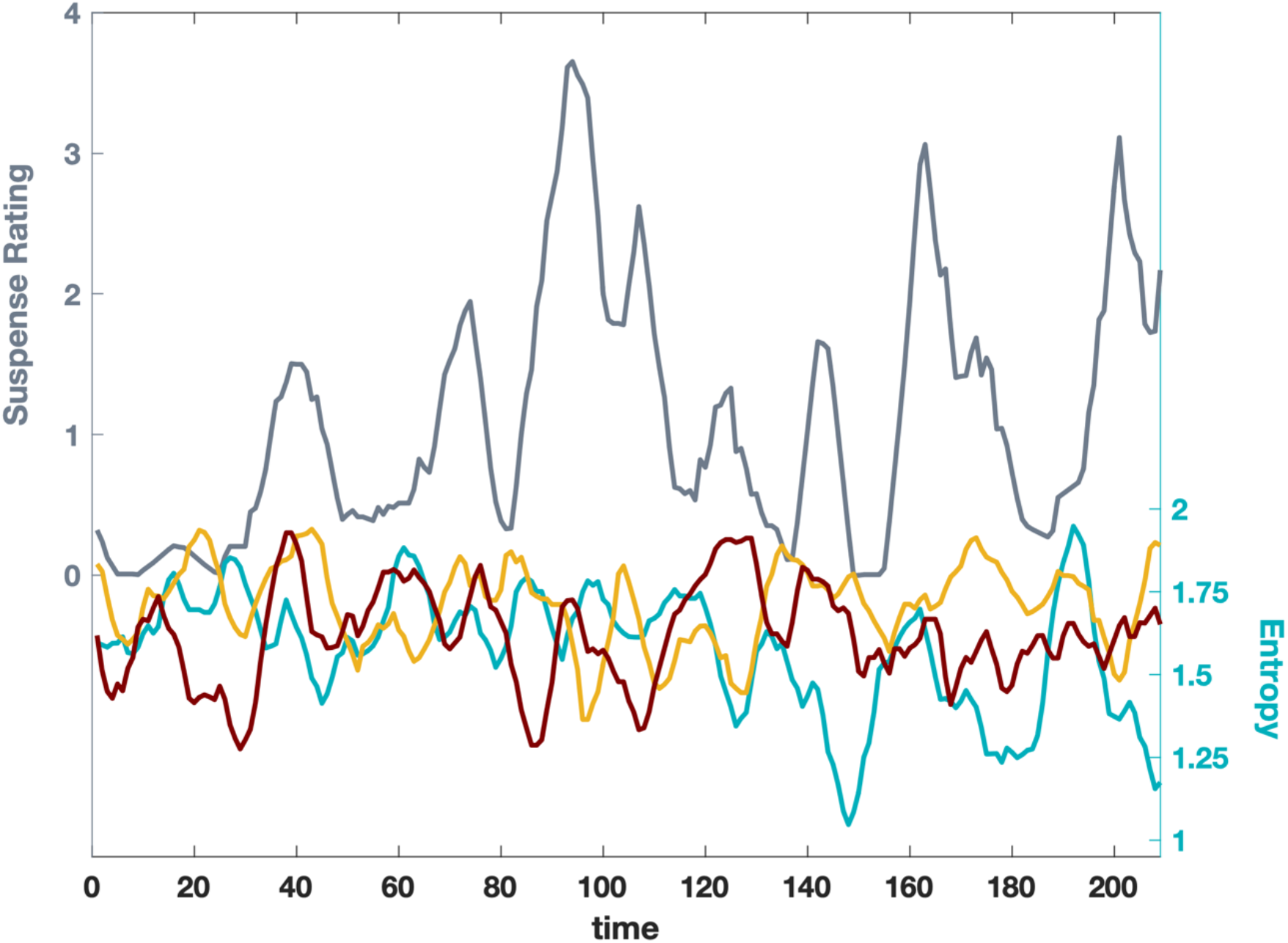
The relationship between the suspense rating and the entropy values in the movie (blue lines), resting state (red lines) and scrambled movie (yellow lines) conditions. Moments with increased suspense was followed by a decrease in the entropy values in the movie watching condition. This negative relationship was not found in the resting state and the scrambled movie watching conditions.

**Table S1.** 42 regions of interest used in the inter-areal coherence analyses. The regions were defined as 10mm-diameter spheres around the given x,y,z coordinates.

| Region of Interest (ROI) | Seed MNI coordinates [x, y, z] |
| --- | --- |
| **Auditory Network (AUD)** |  |
| Anterior cingulate cortex | [6, -7, 43] |
| Precentral gyrus [left] [right] | [-53, -6, 7] [58, -6, 11] |
| Superior transverse temporal gyrus [left] [right] | [-44, -6, 11] [44, -6, 11] |
| **Default Mode Network (DMN)** |  |
| Inferior temporal cortex [left] [right] | [-61, -24, -9] [58, -24, -9] |
| Lateral parietal cortex [left] [right] | [-46, -66, 30] [49, -63, 33] |
| Medial prefrontal cortex | [-1, 54, 27] |
| Posterior cingulate cortex | [0, -52, 27] |
| **Fronto Parietal Network (FP)** |  |
| Angular gyrus [left] [right] | [-31, -59, 42] [30, -61, 39] |
| Midcingulate cortex | [0, -29, 30] |
| Premotor cortex left [left] [right] | [-41, 3, 36] [41, 3, 36] |
| Inferior parietal lobule [left] [right] | [-51, -51, 36] [51, -47, 42] |
| Dorsolateral prefrontal cortex [left] [right] | [-43, 22, 34] [43, 22, 34] |
| **Motor Network (MOT)** |  |
| Supplementary motor area | [0, -21, 48] |
| Primary motor cortex [left] [right] | [-39, -26, 51] [38, -26, 48] |
| **Saliency Network (SAL)** |  |
| Dorsolateral prefrontal cortex [left] [right] | [-38, 52, 10] [30, 48, 22] |
| Ventrolateral prefrontal cortex | [42, 46, 0] |
| Parietal operculum [left] [right] | [-60, -40, 40] [58, -40, 30] |
| Supplementary motor area [left] [right] | [-5, 14, 48] [5, 14, 48] |
| Dorsal anterior cingulate | [-6, 18, 30] |
| Paracingulate cortex | [0, 44, 28] |
| Temporal pole [left] [right] | [-50, 14, -14] [51, 16, -19] |
| Orbital frontoinsula [left] [right] | [-40, 18, -12] [42, 10, -12] |
| **Visual Network (VIS)** |  |
| Associative visual cortex [left] [right] | [30, -89, 20] [-30, -89, 20] |
| Secondary visual cortex [left] [right] | [-6, -78, -3] [6, -78, -3] |
| Primary visual cortex [left] [right] | [-13, -85, 6] [8, -82, 6] |

## REFERENCES

1. Naci, L., Cusack, R., Anello, M. & Owen, A. M. A common neural code for similar conscious experiences in different individuals. Proc. Natl. Acad. Sci. 111, 14277–14282 (2014).

2. Hasson, U., Nir, Y., Levy, I., Fuhrmann, G. & Malach, R. Intersubject Synchronization of Cortical Activity During Natural Vision. Science 303, 1634–1640 (2004).

3. Lankinen, K. et al. Consistency and similarity of MEG- and fMRI-signal time courses during movie viewing. NeuroImage 173, 361–369 (2018).

4. Nummenmaa, L. et al. Emotions promote social interaction by synchronizing brain activity across individuals. Proc. Natl. Acad. Sci. 109, 9599–9604 (2012).

5. Jääskeläinen, I. P. et al. Inter-Subject Synchronization of Prefrontal Cortex Hemodynamic Activity During Natural Viewing. Open Neuroimaging J. 2, 14–19 (2008).

6. Kauppi, J.-P., Jääskeläinen, I., Sams, M. & Tohka, J. Inter-subject correlation of brain hemodynamic responses during watching a movie: localization in space and frequency. Front. Neuroinformatics 4, (2010).

7. Di, X., Zhang, Z., Xu, T. & Biswal, B. B. Dynamic and stable brain connectivity during movie watching as revealed by functional MRI. 2021.09.14.460293 Preprint at https://www.biorxiv.org/content/10.1101/2021.09.14.460293v1 (2021).

8. Meer, J. N. van der, Breakspear, M., Chang, L. J., Sonkusare, S. & Cocchi, L. Movie viewing elicits rich and reliable brain state dynamics. Nat. Commun. 11, 5004 (2020).

9. Hasson, U., Furman, O., Clark, D., Dudai, Y. & Davachi, L. Enhanced Intersubject Correlations during Movie Viewing Correlate with Successful Episodic Encoding. Neuron 57, 452–462 (2008).

10. Simony, E. et al. Dynamic reconfiguration of the default mode network during narrative comprehension. Nat. Commun. 7, 12141 (2016).

11. Damoiseaux, J. S. et al. Consistent resting-state networks across healthy subjects. Proc. Natl. Acad. Sci. 103, 13848–13853 (2006).

12. Cole, M. W., Bassett, D. S., Power, J. D., Braver, T. S. & Petersen, S. E. Intrinsic and Task-Evoked Network Architectures of the Human Brain. Neuron 83, 238–251 (2014).

13. Fox, M. D. et al. The human brain is intrinsically organized into dynamic, anticorrelated functional networks. Proc. Natl. Acad. Sci. 102, 9673–9678 (2005).

14. Laumann, T. O. et al. Functional System and Areal Organization of a Highly Sampled Individual Human Brain. Neuron 87, 657–670 (2015).

15. van den Heuvel, M. P. & Hulshoff Pol, H. E. Exploring the brain network: A review on resting-state fMRI functional connectivity. Eur. Neuropsychopharmacol. 20, 519–534 (2010).

16. Allen, E. A. et al. Tracking whole-brain connectivity dynamics in the resting state. Cereb. Cortex N. Y. N 1991 24, 663–676 (2014).

17. Tagliazucchi, E., Von Wegner, F., Morzelewski, A., Brodbeck, V. & Laufs, H. Dynamic BOLD functional connectivity in humans and its electrophysiological correlates. Front. Hum. Neurosci. 6, (2012).

18. Hutchison, R. M., Womelsdorf, T., Gati, J. S., Everling, S. & Menon, R. S. Resting-state networks show dynamic functional connectivity in awake humans and anesthetized macaques. Hum. Brain Mapp. 34, 2154–2177 (2013).

19. Cabral, J., Kringelbach, M. L. & Deco, G. Functional connectivity dynamically evolves on multiple time-scales over a static structural connectome: Models and mechanisms. NeuroImage 160, 84–96 (2017).

20. Honey, C. J., Kötter, R., Breakspear, M. & Sporns, O. Network structure of cerebral cortex shapes functional connectivity on multiple time scales. Proc. Natl. Acad. Sci. 104, 10240–10245 (2007).

21. Gonzalez-Castillo, J. et al. Tracking ongoing cognition in individuals using brief, whole-brain functional connectivity patterns. Proc. Natl. Acad. Sci. 112, 8762–8767 (2015).

22. Gonzalez-Castillo, J. & Bandettini, P. A. Task-based dynamic functional connectivity: Recent findings and open questions. NeuroImage 180, 526–533 (2018).

23. Barttfeld, P. et al. Signature of consciousness in the dynamics of resting-state brain activity. Proc. Natl. Acad. Sci. U. S. A. 112, 887–892 (2015).

24. Uhrig, L. et al. Resting-state Dynamics as a Cortical Signature of Anesthesia in Monkeys. Anesthesiology 129, 942–958 (2018).

25. Demertzi, A. et al. Human consciousness is supported by dynamic complex patterns of brain signal coordination. Sci. Adv. 5, eaat7603 (2019).

26. Tasserie, J. et al. Deep brain stimulation of the thalamus restores signatures of consciousness in a nonhuman primate model. Sci. Adv. 8, eabl5547.

27. Song, H., Park, B., Park, H. & Shim, W. M. Cognitive and Neural State Dynamics of Narrative Comprehension. J. Neurosci. 41, 8972–8990 (2021).

28. Bezdek, M. A. et al. Neural evidence that suspense narrows attentional focus. Neuroscience 303, 338–345 (2015).

29. Bezdek, M. A., Wenzel, W. G. & Schumacher, E. H. The effect of visual and musical suspense on brain activation and memory during naturalistic viewing. Biol. Psychol. 129, 73–81 (2017).

30. Fox, K. C. R., Spreng, R. N., Ellamil, M., Andrews-Hanna, J. R. & Christoff, K. The wandering brain: Meta-analysis of functional neuroimaging studies of mind-wandering and related spontaneous thought processes. NeuroImage 111, 611–621 (2015).

31. Mason, M. F. et al. Wandering minds: the default network and stimulus-independent thought. Science 315, 393–395 (2007).

32. Haugg, A. et al. Do Patients Thought to Lack Consciousness Retain the Capacity for Internal as Well as External Awareness? Front. Neurol. 9, (2018).

33. Vanhaudenhuyse, A. et al. Two distinct neuronal networks mediate the awareness of environment and of self. J. Cogn. Neurosci. 23, 570–578 (2011).

34. Schiff, N. D. Cognitive Motor Dissociation Following Severe Brain Injuries. JAMA Neurol. 72, 1413–1415 (2015).

35. Fernández-Espejo, D. & Owen, A. M. Detecting awareness after severe brain injury. Nat. Rev. Neurosci. 14, 801–809 (2013).

36. Whitfield-Gabrieli, S. & Nieto-Castanon, A. Conn: A functional connectivity toolbox for correlated and anticorrelated brain networks. Brain Connect. 2(3), 125–141 (2012).

37. Bates, D., Mächler, M., Bolker, B. & Walker, S. Fitting Linear Mixed-Effects Models Using lme4. J. Stat. Softw. 67, 1–48 (2015).

38. Fox, J. & Weisberg, S. An R Companion to Applied Regression. (Sage, 2019).

39. Lenth, R. V. emmeans: Estimated Marginal Means, aka Least-Squares Means. R package version 1.6.2-1. (2021).

40. R Core Team. R: A Language and Environment for Statistical Computing. (R Foundation for Statistical Computing, 2021).

